# A clinically-relevant polymorphism in the Na^+^/taurocholate cotransporting polypeptide (NTCP) occurs at a rheostat position

**DOI:** 10.1101/2020.07.02.184424

**Authors:** Melissa J. Ruggiero, Shipra Malhotra, Aron W. Fenton, Liskin Swint-Kruse, John Karanicolas, Bruno Hagenbuch

## Abstract

Conventionally, most amino acid substitutions at important protein positions are expected to abolish function. However, in several soluble-globular proteins, we identified a class of non-conserved positions for which various substitutions produced progressive functional changes; we consider these evolutionary “rheostats”. Here, we report a strong rheostat position in the integral membrane protein, Na^+^/taurocholate cotransporting polypeptide (NTCP), at the site of a pharmacologically-relevant polymorphism (S267F). Functional studies were performed for all 20 substitutions (“S267X”) with three substrates (taurocholate, estrone-3-sulfate and rosuvastatin). The S267X set showed strong rheostatic effects on overall transport, and individual substitutions showed varied effects on transport kinetics (K_m_ and V_max_). However, the outcomes were substrate dependent, indicating altered specificity. To assess protein stability, we measured surface expression and used the Rosetta software suite to model structure and stability changes of S267X. Although buried near the substrate binding site, S267X substitutions were easily accommodated in the NTCP structure model. Across the modest range of changes, calculated stabilities correlated with surface-expression differences, but neither parameter correlated with altered transport. Thus, substitutions at rheostat position 267 had wide-ranging effects on the phenotype of this integral membrane protein. We further propose that polymorphic positions in other proteins might be locations of rheostat positions.

## Introduction

Amino acid substitutions are commonly used to evaluate which amino acids in a protein contribute to function. Several decades of studies have led to conventional “rules” for mutational outcomes that are now included in many textbooks and are often implicitly or explicitly assumed in the design and interpretation of experimental studies. For instance, at “important” protein positions, only amino acids with biochemical properties similar to the wild-type are expected to allow function, whereas other amino acid substitutions are expected to abolish function or structure. However, the mutational studies that gave rise to these rules were primarily focused on evolutionarily conserved amino acid positions (Gray, Kukurba, & Kumar, 2012). When we performed substitution studies of less conserved positions, results were seldom consistent with expected outcomes. Instead of an “on/off” pattern, when non-conserved positions were substituted with a variety of amino acids, each substitution had a different outcome (*e.g*. Figure 2). The fact that one position could be substituted to access a continuum of functional outcomes is analogous to an electronic dimmer switch; therefore, these positions have been labeled as “rheostat” positions (Hodges, Fenton, Dougherty, Overholt, & Swint-Kruse, 2018; Meinhardt, Manley, Parente, & Swint-Kruse, 2013; Wu, Swint-Kruse, & Fenton, 2019).

At the onset of the current study, we had three areas of interest. First and foremost, we were curious whether rheostat positions were limited to the soluble-globular class of proteins in which they were discovered, or if they also exist in transmembrane proteins (Meinhardt et al., 2013). Second, if rheostat positions do exist in integral membrane proteins, we wondered whether the functional outcomes arising from various substitutions were dependent upon the substrate being transported. That is, we wished to explore the effects of substitutions at rheostat positions on substrate specificity. Third, we wanted to relate the continuum of functional outcomes to the complex conformational changes experienced by an integral membrane protein during transport.

To address these questions, we chose to use human Na^+^/taurocholate cotransporting polypeptide (NTCP) as a model membrane protein. NTCP is expressed at the basolateral membrane of human hepatocytes where it plays an important role in the enterohepatic circulation of bile acids (Hagenbuch & Dawson, 2004). In addition to conjugated bile acids such as taurocholate (TCA), NTCP mediates the uptake of other substrates into hepatocytes, including estrone-3-sulfate and several statins such as rosuvastatin (Claro da Silva, Polli, & Swaan, 2013). Furthermore, several single nucleotide polymorphisms (SNPs) have been reported to alter the transport activity of NTCP (Ho, Leake, Roberts, Lee, & Kim, 2004). One of these SNPs (NTCP*2) leads to the missense amino acid substitution S267F and has an allele frequency of 7.5% in Chinese Americans. Previously published data for S267F indicated reduced transport of taurocholate, wild-type-like transport of estrone-3-sulfate, and increased transport of rosuvastatin *in vitro* (Ho et al., 2004; Ho et al., 2006; Pan et al., 2011). Clinically, this mutation resulted in severe hypercholanemia with total serum bile acid levels of about 15- to 70-fold above normal in homozygous pediatric patients (Deng et al., 2016; Dong et al., 2019); some of the patients also had elevated liver enzymes, jaundice and gallstones (Dong et al., 2019). In homozygous adult patients, NTCP*2 resulted in total serum bile acid levels 2- to 5-fold above normal (Liu et al., 2017).

We interpreted these previous observations as a potential indication that NTCP bile acid transport could be rheostatically modulated by substitutions at position 267. To test this, we assessed the function of wild-type NTCP and all 19 amino acid substitutions at position 267 with cellular uptake studies. We also determined whether substitution outcomes were substrate dependent, by measuring transport of taurocholate, estrone-3-sulfate, and rosuvastatin. Additional experiments aimed to differentiate the effects of substitutions on protein surface expression and transport kinetics. Finally, we used homology modeling and energetic calculations to relate the rheostatic modulation of function to the complex conformational changes utilized by NTCP to transport bile acids and other substrates (Zhou et al., 2014).

## Materials and Methods

### Materials

Radiolabeled [^3^H]-taurocholate (6.5 Ci/mmol) was purchased from PerkinElmer (Boston, MA). [^3^H]-Estrone-3-sulfate (50 Ci/mmol) and [^3^H]-rosuvastatin (10 Ci/mmol) were from American Radiolabeled Chemicals (St. Louis, MO). Taurocholic acid sodium salt (97% pure) and estrone-3-sulfate sodium salt (containing 35% Tris stabilizer) were purchased from Sigma Aldrich (St. Louis, MO). Rosuvastatin (98% pure) was purchased from Cayman Chemicals (Ann Arbor, Michigan).

### Site-directed mutagenesis

Mutagenesis reactions were completed using the QuikChange Lightning Multi Site-Directed Mutagenesis Kit (Agilent, Santa Clara, CA). We used a previously-cloned His-tagged human NTCP in the pcDNA5/FRT vector for the mutagenesis template (Zhao et al., 2015). Primers were designed to flank the S267 codon by approximately twenty nucleotides and were ordered from Invitrogen (Carlsbad, CA). Mutated cDNA was then transformed into One Shot TOP10 cells (Thermo Fisher Scientific, Waltham, MA) and bacterial colonies were randomly selected. The cDNA was isolated using QIAGEN (Hilden, Germany) mini prep kits and sequenced (GENEWIZ, South Plainfield, NJ). Constructs containing the appropriate amino acid replacements were transfected into HEK293 cells for functional or expression studies as described below.

### Cell culture

HEK293T/17 cells obtained from (American Type Culture Collection, Manassas, VA) were plated on poly-D-lysine coated flat bottom 48- and 6-well TPP cell culture plates at densities of 100,000 and 800,000 cells per well respectively. Cells were cultured in Dulbecco’s Modified Eagle’s Medium (American Type Culture Collection) containing 100 U/ml penicillin, 100 μg/ml streptomycin (Thermo Fisher Scientific) and 10% fetal bovine serum (Hyclone, Thermo Fisher Scientific) for 24 hours prior to transfection. Experiments were then completed 48 hours post transfection.

### Transient transfection of HEK293 cells

Empty vector, wild-type NTCP, and variant NTCP plasmids were transfected into HEK293 cells 24 hours after plating. Transfections followed the FuGENE HD protocol obtained from Promega (Madison, WI). All plasmids were transfected in triplicates for functional studies on 48-well plates. For surface biotinylation experiments completed on 6-well plates, single wells were transfected.

### Initial uptake experiments

The uptake procedure has been optimized and is well established in our laboratory for functional studies (Zhao et al., 2015). Uptake buffer (142 mM NaCl, 5 mM KCl, 1 mM KH_2_PO_4_, 1.2 mM MgSO_4_, 1.5 mM CaCl_2_, 5 mM glucose, and 12.5 mM HEPES, pH 7.4) was used for all washes and to prepare uptake solutions. Cells were washed with warm uptake buffer and then incubated at 37°C with nanomolar concentrations of either [^3^H]-taurocholate (30 nM), [^3^H]-estrone-3-sulfate (5.8 nM), or [^3^H]-rosuvastatin (50 nM) for five minutes. Uptake was then terminated by washing with ice-cold uptake buffer and the cells were solubilized in a 1% TX-100 solution in phosphate buffered saline (PBS). Radioactivity was measured using a liquid scintillation counter and protein concentrations were determined using bicinchoninic acid assays (Thermo Fisher Scientific). Results were calculated by correcting for total protein, subtracting the uptake by cells expressing empty vector, setting wild-type NTCP as 100%, and then comparing all variants to wild-type as percent of control.

### Surface biotinylation

HEK293 cells were plated on 6-well plates and transfected as described above. Forty-eight hours later, cells were incubated for 15 minutes on ice, and all solutions and buffers used were prechilled. Each well was washed with PBS and then incubated for one hour at 4°C while rocking with 1 mg/mL EZ-Link NHS-SS-Biotin (Thermo Fisher Scientific) in PBS. Cells were then washed with PBS and incubated for twenty minutes at 4°C with 100 mM glycine in PBS while rocking. After washing with PBS, cells were lysed using 10 mM Tris, 150 mM NaCl, 1 mM EDTA, 0.1% SDS, 1% Triton X-100, pH 7.5 (lysis buffer), containing protease inhibitors (cOmplete protease inhibitor cocktail, Sigma-Aldrich). The lysates were centrifuged at 10,000 x g for two minutes and the supernatant was incubated for one hour at room temperature using end-over-end rotation with pre-washed NeutrAvidin Agarose Resin beads (Thermo Fisher Scientific). The beads were washed with lysis buffer, captured proteins were eluted with 1 X SDS sample buffer containing 5% β-mercaptoethanol and 1 X protease inhibitors at 70°C for 10 minutes and collected by centrifugation at 850 x g for five minutes.

### Western blotting

Surface biotinylation samples were heated to 50°C for 10 minutes before separation using 4-20% polyacrylamide gradient gels (Bio-Rad, Hercules, CA). After separation, proteins were transferred to nitrocellulose membranes using Invitrogen’s Power Blotter System. Blots were blocked with 5% milk in Tris-buffered saline containing 0.1% Tween 20 (TBS-T) for one hour at room temperature while rocking. Following blocking, blots were incubated overnight at 4°C with a mouse antibody against the α subunit of Na^+^/K^+^-ATPase (Abcam-ab7671, 1:2,000) and with a mouse antibody against the His-tag (Tetra·His Antibody, QIAGEN-Cat. No.34670, 1:2,000) in blocking solution on a rocker. The next day blots were washed with TBS-T and TBS before incubation with an HRP conjugated goat anti-mouse secondary antibody at 1:10,000 (Thermo Fisher Scientific-Cat. No. 31430) in 2.5% milk in TBS. After one hour, blots were washed with TBS and incubated with SuperSignal West Pico Chemiluminescent substrate (Thermo Fisher Scientific). Blots were visualized using a LI-COR Odyssey Fc (LI-COR, Lincoln, NE) and bands were quantified using their Image Studio Lite Quantification Software.

### Time dependency and kinetics experiments

Initial linear rates were determined for wild-type, S267F, S267W, and S267N using multiple time points between 10 seconds and 10 minutes at low and high concentrations of each substrate: taurocholate at 0.1 μM and 100 μM; estrone-3-sulfate at 1 μM and 200 μM; and rosuvastatin at 5 μM and 500 μM (Table 1). Kinetics for wild-type, S267F, S267W, and S267N were determined using HEK293 cells plated on 48-well plates. Uptakes were performed using sodium containing and sodium-free uptake buffers (NaCl was replace by choline chloride) and after correction for total protein and surface expression, results were analyzed using GraphPad Prism 8 (Michaelis-Menten kinetics).

**Table 1.**
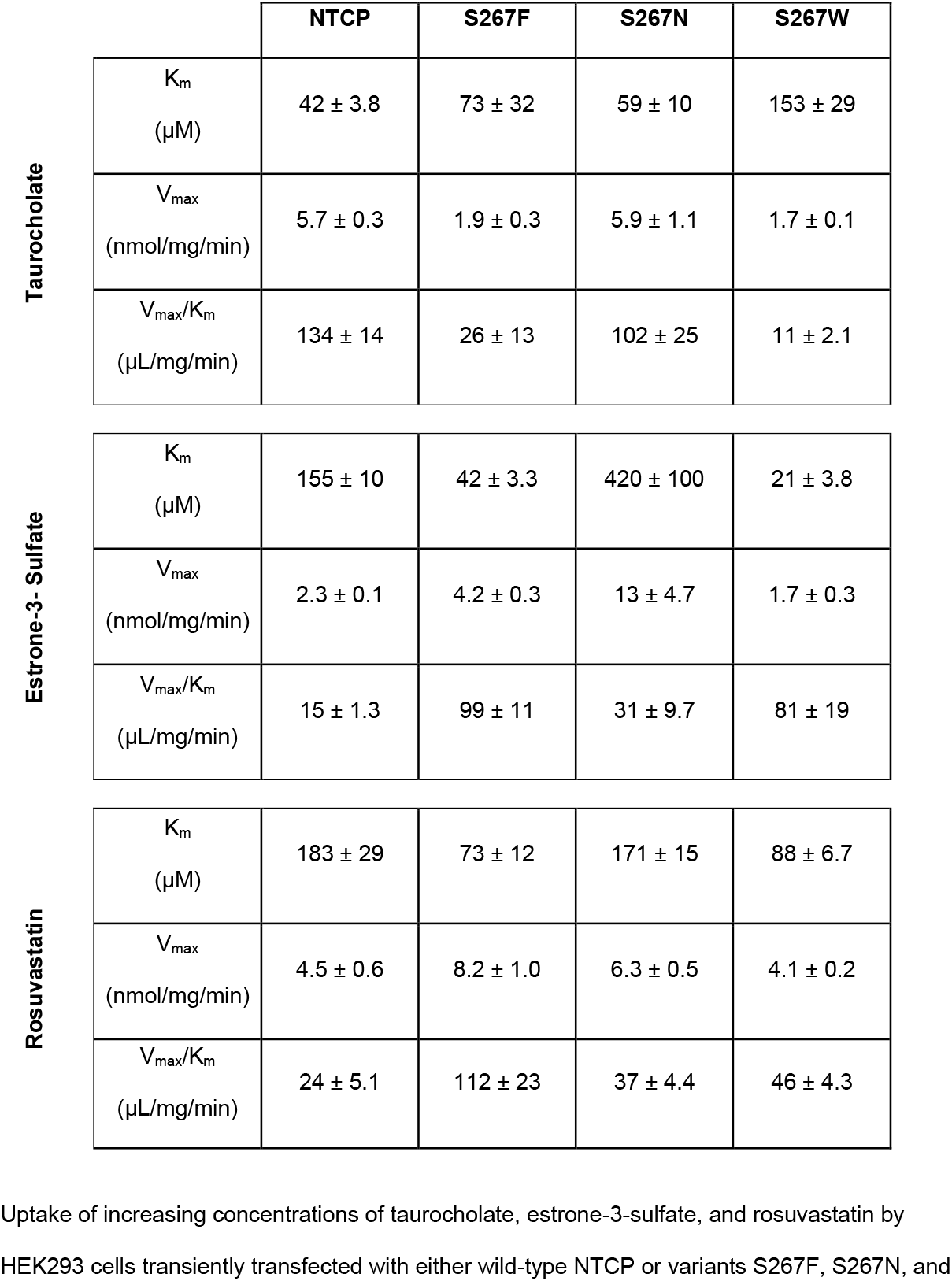
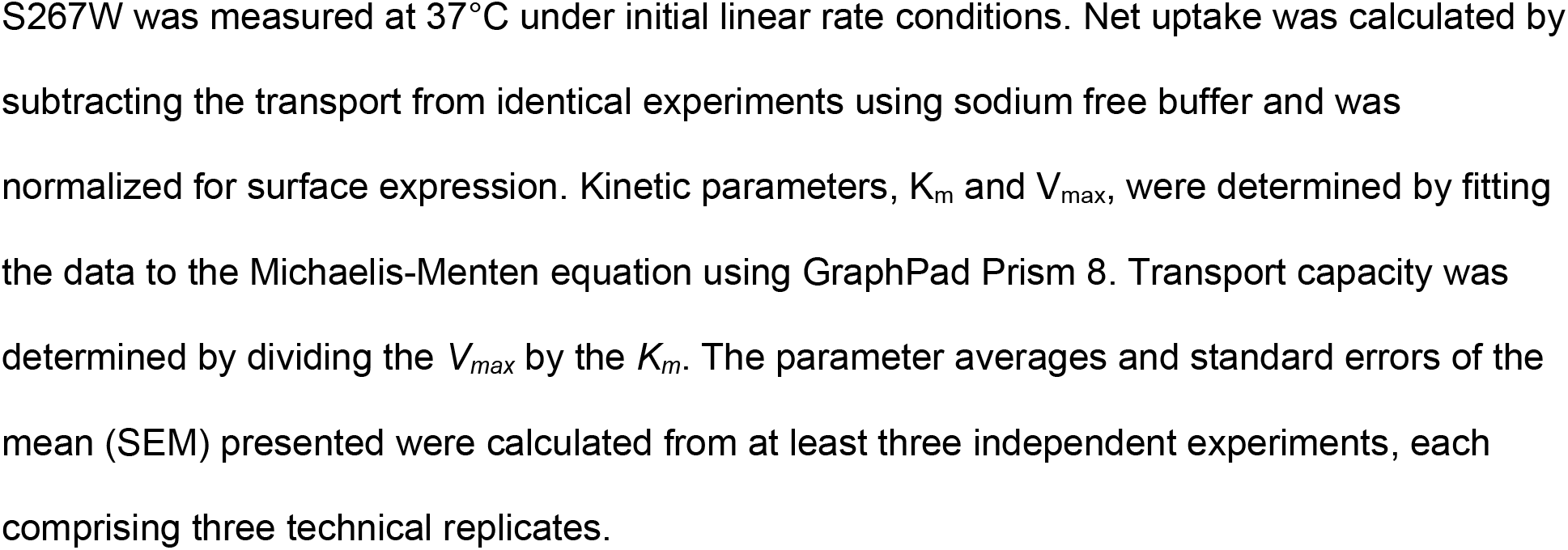
Kinetic values for substrate uptake mediated by wild-type NTCP and selected variants

|  |  | **NTCP** | **S267F** | **S267N** | **S267W** |
| --- | --- | --- | --- | --- | --- |
| **Taurocholate** | $K_m$<br>($\mu\text{M}$) | $42 \pm 3.8$ | $73 \pm 32$ | $59 \pm 10$ | $153 \pm 29$ |
| | $V_{\max}$<br>(nmol/mg/min) | $5.7 \pm 0.3$ | $1.9 \pm 0.3$ | $5.9 \pm 1.1$ | $1.7 \pm 0.1$ |
| | $V_{\max}/K_m$<br>($\mu\text{L}/\text{mg}/\text{min}$) | $134 \pm 14$ | $26 \pm 13$ | $102 \pm 25$ | $11 \pm 2.1$ |
| **Estrone-3- Sulfate** | $K_m$<br>($\mu\text{M}$) | $155 \pm 10$ | $42 \pm 3.3$ | $420 \pm 100$ | $21 \pm 3.8$ |
| | $V_{\max}$<br>(nmol/mg/min) | $2.3 \pm 0.1$ | $4.2 \pm 0.3$ | $13 \pm 4.7$ | $1.7 \pm 0.3$ |
| | $V_{\max}/K_m$<br>($\mu\text{L}/\text{mg}/\text{min}$) | $15 \pm 1.3$ | $99 \pm 11$ | $31 \pm 9.7$ | $81 \pm 19$ |
| **Rosuvastatin** | $K_m$<br>($\mu\text{M}$) | $183 \pm 29$ | $73 \pm 12$ | $171 \pm 15$ | $88 \pm 6.7$ |
| | $V_{\max}$<br>(nmol/mg/min) | $4.5 \pm 0.6$ | $8.2 \pm 1.0$ | $6.3 \pm 0.5$ | $4.1 \pm 0.2$ |
| | $V_{\max}/K_m$<br>($\mu\text{L}/\text{mg}/\text{min}$) | $24 \pm 5.1$ | $112 \pm 23$ | $37 \pm 4.4$ | $46 \pm 4.3$ |

Uptake of increasing concentrations of taurocholate, estrone-3-sulfate, and rosuvastatin by HEK293 cells transiently transfected with either wild-type NTCP or variants S267F, S267N, and
S267W was measured at 37°C under initial linear rate conditions. Net uptake was calculated by subtracting the transport from identical experiments using sodium free buffer and was normalized for surface expression. Kinetic parameters, $K_m$ and $V_{max}$ , were determined by fitting the data to the Michaelis-Menten equation using GraphPad Prism 8. Transport capacity was determined by dividing the $V_{max}$ by the $K_m$ . The parameter averages and standard errors of the mean (SEM) presented were calculated from at least three independent experiments, each comprising three technical replicates.

### Homology model for human NTCP

To model the human NTCP sequence (NCBI: NP_003040), structural models were constructed using the SWISS-MODEL automated protein modeling server (https://swissmodel.expasy.org/) (Arnold, Bordoli, Kopp, & Schwede, 2006; Benkert, Biasini, & Schwede, 2011; Biasini et al., 2014). To model the inward-open conformation of NTCP, we used as a template the structure of a bacterial homologue from *Neisseria meningitidis* (PDB 3ZUY, 25% sequence similarity with NTCP) (Hu, Iwata, Cameron, & Drew, 2011). To model the outward-open conformation of NTCP, we used as a template the structure of a bacterial homologue from *Yersinia frederiksenii* (PDB 4N7X, 26% sequence similarity with NTCP) (Zhou et al., 2014). As with the templates upon which the models were based, the topology of each model comprises 9 transmembrane (TM) helices linked by short loops into core (TM 3,4,5,8,9 and 10) and panel domains (TM 2,6 and 7), along with the substrate binding intracellular crevice (Figure 1). The model of NTCP lacks the N-terminal sequence corresponding to TM1 of the ASBT crystal structures; to keep the helix numbering consistent with the known crystal structures, we have chosen to start the NTCP helix numbering with TM2.

**Figure 1:**
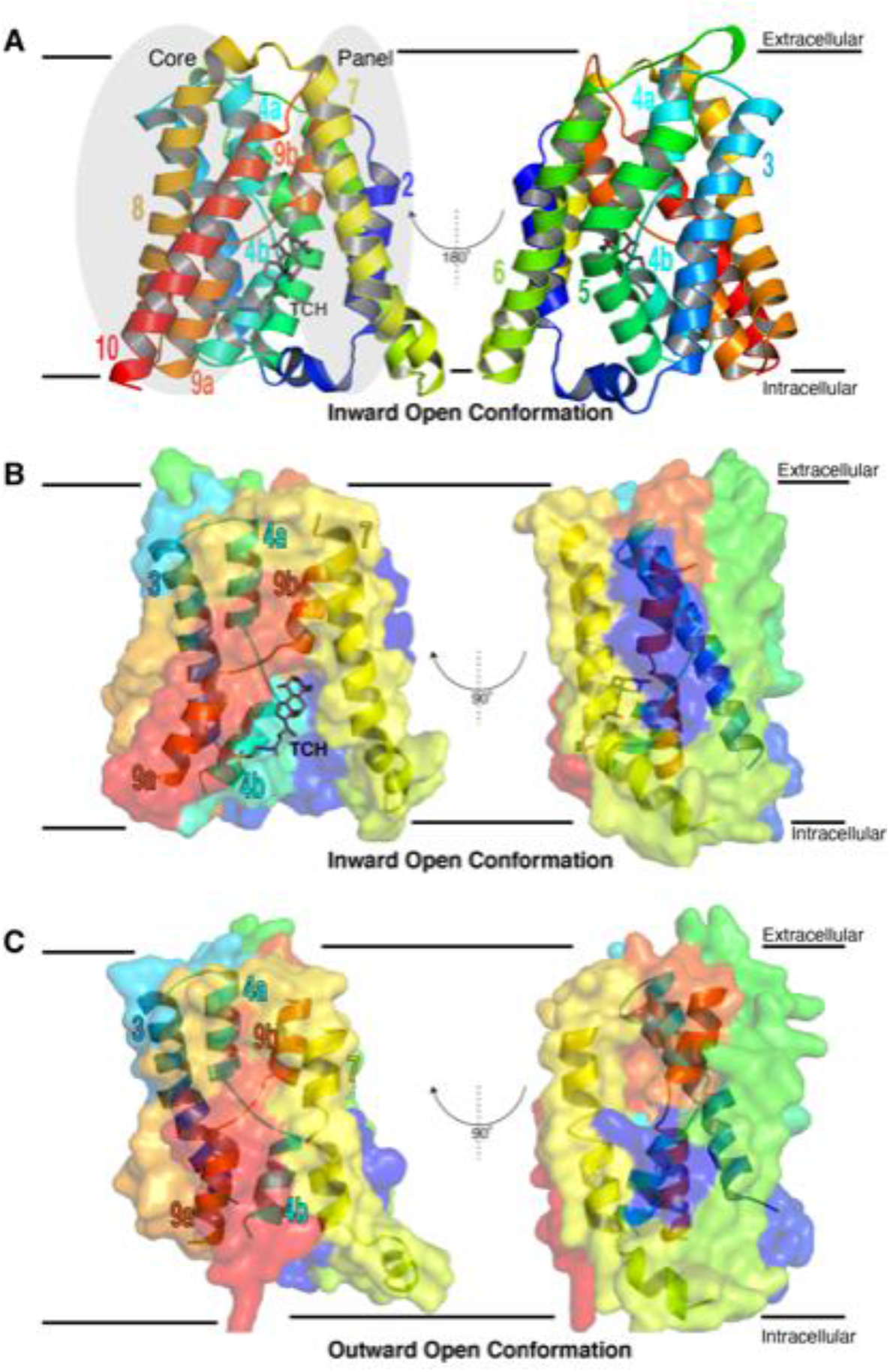
Comparative models of human NTCP. (A) Homology model of human NTCP in the inward-open conformation, built using ASBT_NM_ as a structural template. The structure comprises 9 TM helices (denoted TMs 2, 3, 4a, 4b, 5-8, 9a, 9b, and 10). Position 267 is located on TM 9b. The inward-open conformation has a large crevice at the intracellular side of membrane that is formed between the core and panel domains. Taurocholate, in gray sticks, is included in this perspective by superimposition from the template to show the interaction of a substrate in the inward open conformation. (B) The inward-open model shown in cross-section, to highlight the arrangement of helices in this conformation. (C) Homology model of human NTCP in the outward-open conformation, built using ASBT_YF_ as a structural template. Relative to the inward-open conformation, the substrate-binding pocket has closed in this conformation, due to a concerted movement of TM helices 3, 7, 4a, 9b, 4b, and 9a.

### Modeling S267 variants of human NTCP

To facilitate structural exploration in response to sequence variants, we began by using the “relax” protocol (Conway, Tyka, DiMaio, Konerding, & Baker, 2014; Nivon, Moretti, & Baker, 2013; Tyka et al., 2011) in the Rosetta macromolecular modeling suite (Leaver-Fay et al., 2011) to generate a close structural ensemble from each of the two homology models provided by SWISS-MODEL. Each starting conformation was used to carry out 1000 independent simulations and the top-scoring 100 output structures were retained as a representative ensemble for the (wild-type) inward-open or outward-open state.

To build a structural model of a given NTCP sequence variant, we used the “ddG” protocol in Rosetta (Kellogg, Leaver-Fay, & Baker, 2011; Kuhlman et al., 2003). With respect to our study, this protocol was used to introduce the desired amino acid substitution at position 267 and then iterated between optimization of the nearby sidechains and optimization of the backbone. We applied this protocol 10 times to each of the 100 members of our (wild-type) structural ensemble to yield 1000 models of the desired sequence variant. To avoid potential sampling artifacts from drawing the conformation/energy from the single lowest-energy conformation sampled, we ranked all conformations for a given sequence variant on the basis for Rosetta energy and carried forward the 50^th^-best conformation (*i.e*., 95^th^ percentile) as the representative.

The same process was repeated for each of the 20 amino acids, to generate all possible sequence variants at this position. While the starting models already included serine at position 267, the same protocol was nonetheless applied to introduce serine; this ensured that any structural/energetic changes were indeed due to sequence variations and not simply changes relative to the starting structure induced by the modeling protocol.

The same analysis was separately completed using the structural ensemble for the inward-open state and the structural ensemble for the outward-open state.

All Rosetta calculations were carried out using git revision 0e7ed9fd3cd610f2a7c9f3bdcaba64a9b11aab0d of the developer master source code.

### Statistical analysis

Calculations were performed using GraphPad Prism 8 (GraphPad Software Inc., San Diego, CA). Correlation was evaluated using Pearson and Spearman correlation coefficients. Significance was determined using One-way ANOVA followed by Dunnett’s multiple comparisons. Results were considered significantly different at p < 0.05.

### Construction of an NTCP sequence alignment

A detailed description of the NTCP sequence alignment is given in the Supplemental Methods. The references associated with these methods are included in the Supplemental Material.

## Results

### Cellular substrate transport of S267 variants

A property of the rheostat positions previously observed in soluble-globular proteins is that they were non-conserved throughout evolution (*e.g*. (Meinhardt et al., 2013)). NTCP position 267 was reported to be highly conserved among many different animals, including: primates, rodents, dogs, cats, horses, chicken, several fish, and marine chordate (Deng et al., 2016). However, when we expanded the sequence alignment to include 1,561 homologues of the Solute Carrier Family 10A (SLC10A) (Claro da Silva et al., 2013) representing all kingdoms of life (see sequence alignment file in the Supplemental Material), position 267 had a sequence entropy of 1.54. This intermediate conservation score (overall range of 0.0 to 2.8) suggested that position 267 could tolerate multiple substitutions without catastrophic outcomes. This type of outcome is expected to occur for several amino acid replacements at a rheostat position and is consistent with the documented non-catastrophic (and indeed functional) changes for the known polymorphism at position 267 in NTCP, as noted in the introduction.

To ascertain whether position 267 was a rheostat position, we replaced serine with all other nineteen amino acids and measured uptake of taurocholate (Figure 2, top), estrone-3-sulfate (Figure 2, middle), and rosuvastatin (Figure 2, bottom) for each substitution. On the left side of Figure 2, uptake is shown with the amino acids in alphabetical order, with wild-type at far left. The right panels show results ordered from highest to lowest transport, with wild-type placed within the series.

**Figure 2:**
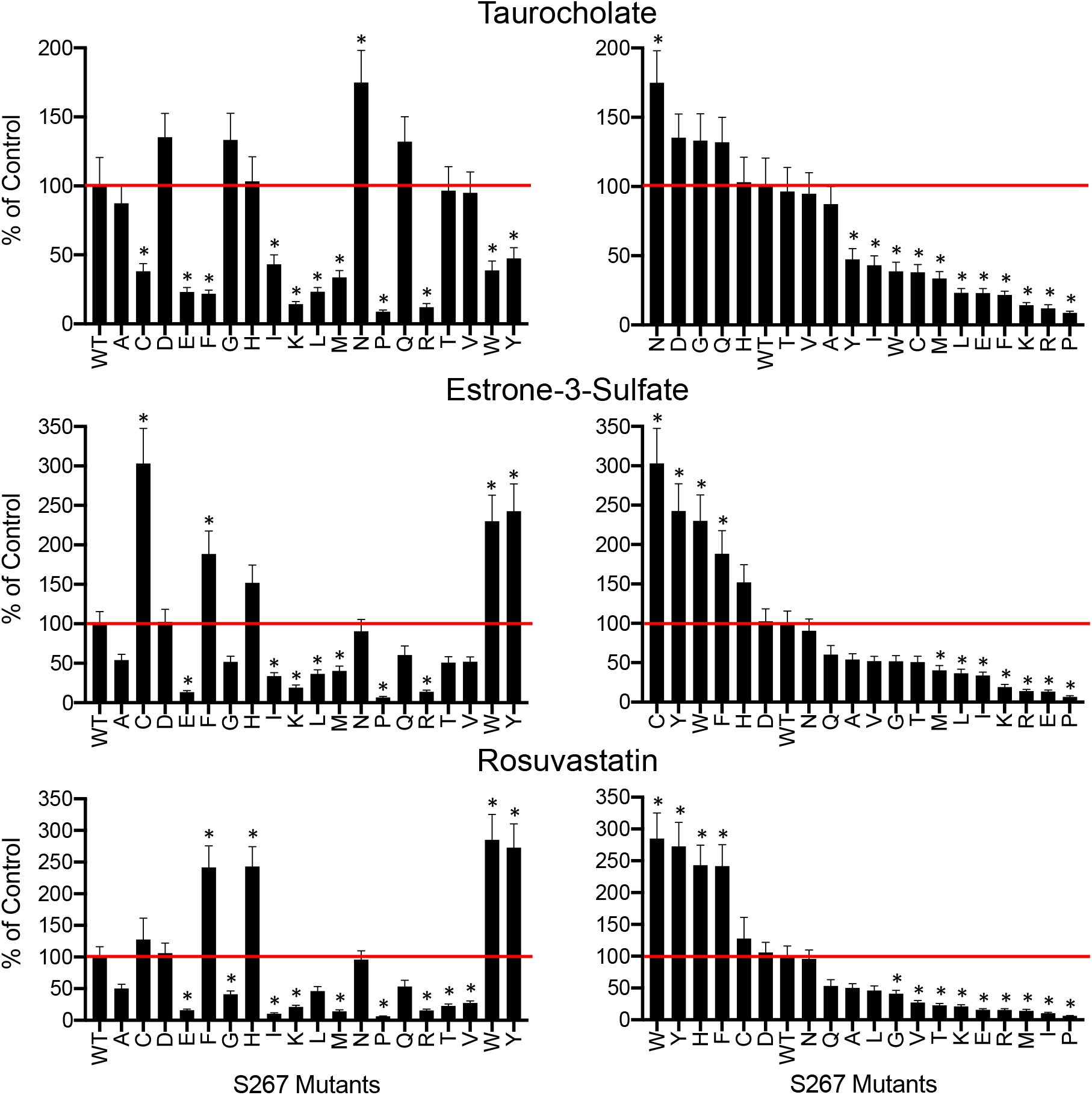
Substrate uptake by wild-type NTCP and S267 variants. Uptake of ^3^[H]taurocholate (30 nM), ^3^[H]estrone-3-sulfate (5.8 nM) and ^3^[H]rosuvastatin (50 nM) was measured for 5 min at 37°C, 48 hours after transfection of wild-type NTCP (WT), its S267 variants and empty vector into HEK293 cells. Net uptake was obtained by subtracting the uptake of cells transfected with empty vector from uptake of NTCP-expressing cells. The left-hand side shows the results ordered alphabetically based on the amino acid replacement and the right-hand side shows the substitutions ordered from highest to lowest transport activity. Results were calculated as percent of wild-type NTCP and are reported as the mean ± SEM from at least three independent experiments, each comprising three technical replicates. Horizontal lines to aid visual inspection correspond to wild-type (“WT”) values, which were set to 100%, and asterisks indicate a p < 0.05 level of significant difference from wild-type NTCP.

Notably, for all three substrates, some variants transported substrates better than wild-type whereas others had diminished transport. The range of observed changes spanned several orders of magnitude. Thus, position 267 exhibited definitive “rheostatic” substitution behavior. Furthermore, the rank-order of the amino acid substitutions differed among the three substrates. This is further discussed below, but here we particularly note that the NTCP*2 polymorphism, S267F, showed a significant decrease in taurocholate transport, a 2.0-fold increase for estrone-3-sulfate transport, and 2.5-fold increase for rosuvastatin transport. Previous reports showed decreased taurocholate transport, estrone-3-sulfate levels similar to wild-type, and increased rosuvastatin transport (Ho et al., 2004; Ho et al., 2006; Pan et al., 2011). The discrepancy for estrone-3-sulfate could arise from the slightly different uptake conditions including substrate concentrations, incubation time, and/or different cell lines used in the two studies.

### Dissecting the composite cellular outcomes of S267 variants

Protein substitutions can alter substrate transport kinetics, substrate specificity, protein stability, and/or intra-cellular trafficking to the outer membrane. The cellular uptake assay is sensitive to changes in any of these parameters. Thus, we devised experiments to dissect the functional and structural contributions.

To assess the combined effects of trafficking and stability, we quantified differences in surface expression of NTCP variants using surface biotinylation experiments followed by western blotting (Figure 3A). When normalized for the loading control, Na^+^/K^+^ATPase, the expression levels varied between 22.5% for S267Q and 153% for S267W. The expression of most of the other variants was similar to wildtype (Figure 3B). Thus, NTCP appeared to accommodate different amino acid side chains at position 267 without significantly disrupting the overall structure or trafficking to the cell surface.

**Figure 3.**
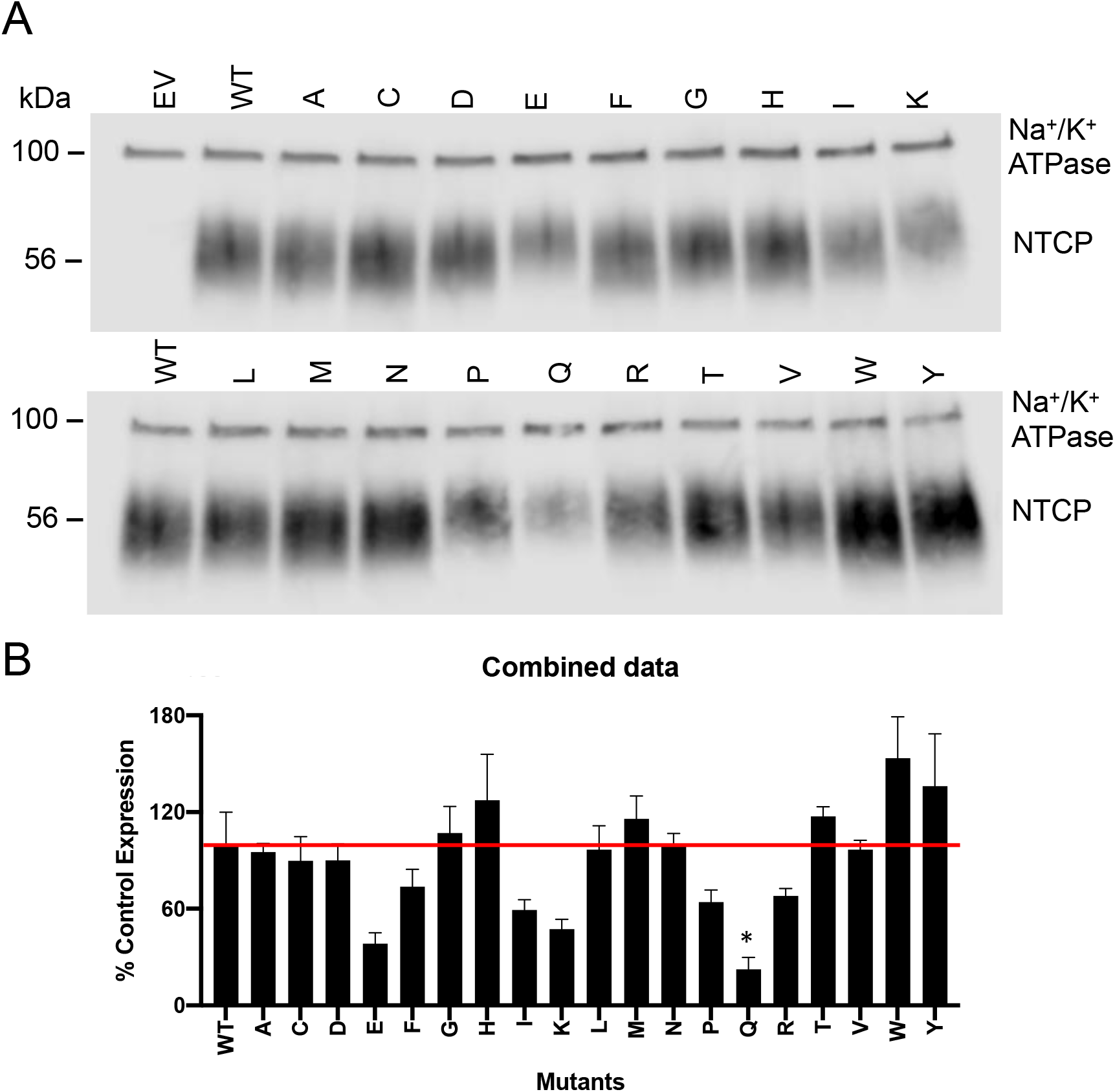
Surface expression of wild-type NTCP and S267 variants. (A) Representative western blot of surface expressed wild-type NTCP and its S267 variants in transiently transfected HEK293 cells. Empty vector (EV), wild-type (WT), and S267 variant proteins were separated on a 4-20% gel and then transferred to a nitrocellulose membrane. Blots were probed with a combination of Na^+^/K^+^-ATPase (loading control at 100 kDa) and tetra-His antibodies (recognizes the His-tagged transporter). (B) Quantification of S267 variants relative to wild-type NTCP. Expression was quantified using Image Studio Lite and the bars represent the mean ± SEM of three independent experiments; an asterisk indicates a p < 0.05 level of significant difference from wild-type NTCP. The horizontal line indicates wild-type control, which was set to 100%.

Next, initial uptake experiments from Figure 2 were corrected for the surface expression and results are shown in Figure 4. The overall rheostat-like behavior remained, showing that substitutions altered transport function. The rank orders of S267 substitutions were further analyzed using correlation plots to illustrate the effects on substrate specificity (Figure 5, Supplemental Table 1). Strikingly, taurocholate uptake did not correlate with either estrone-3-sufate or rosuvastatin uptake (Figure 5A and B). Because the outcome of each substitution differed among the substrates, these data indicate that amino acid substitutions at rheostat position 267 altered substrate specificity (Tungtur, Schwingen, Riepe, Weeramange, & Swint-Kruse, 2019). In contrast, with the exception of S267C, the uptake of estrone-3-sulfate and rosuvastatin strongly correlated (Figure 5C), suggesting a shared translocation pathway for these two substrates.

**Figure 4.**
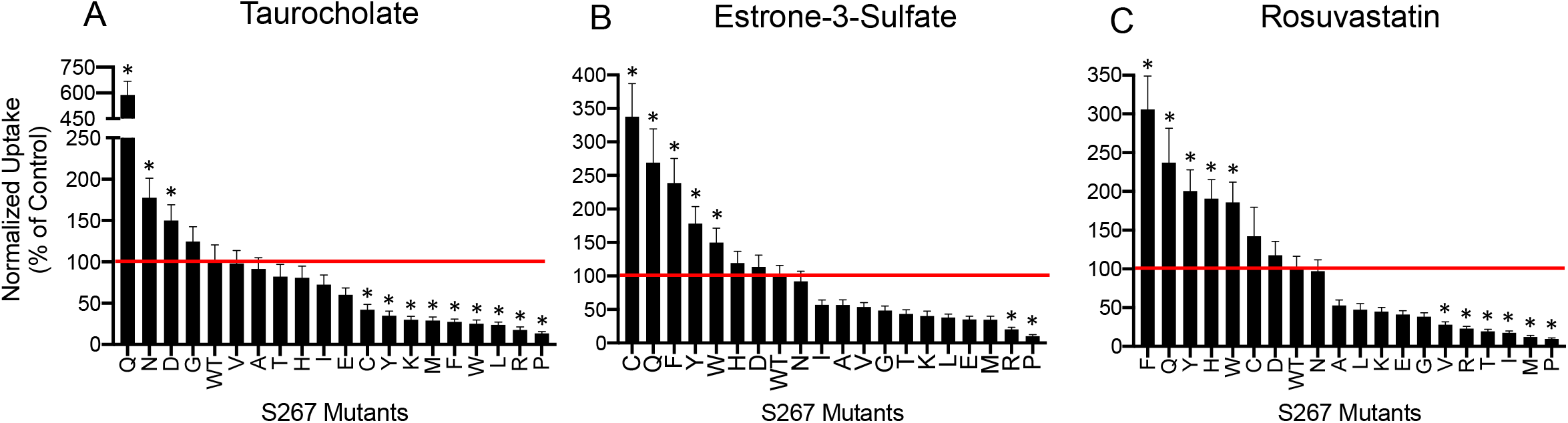
Initial substrate uptake normalized for surface expression. Uptake results (Figure 1) were corrected for the surface expression (Figure 2B) and are presented with the substitutions rank-ordered from largest to smallest for each substrate. Horizontal lines indicate wild-type control, which was set to 100%. Error bars represents propagated SEM; asterisks indicate a p < 0.05 level of significant difference from wild-type NTCP.

**Figure 5.**
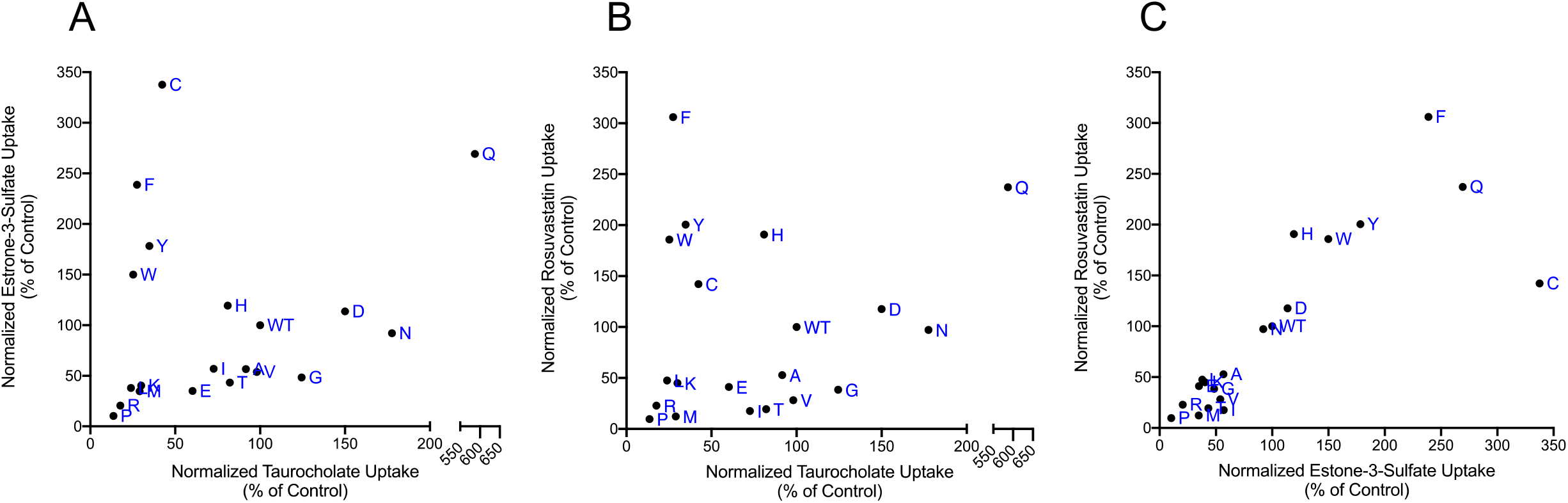
Comparison of normalized substrate uptake among the three different substrates. Normalized uptake values from Figure 3 are plotted against each other. (A) ^3^[H]taurocholate (TCA) versus ^3^[H]estrone-3-sulfate (E3S), (B) TCA versus ^3^[H]rosuvastatin, and (C) E3S versus ^3^[H]rosuvastatin. For E3S versus rosuvastatin, the Pearson Coefficient, which is a measure of the linear correlation, was 0.82 (p < 0.0001) and the Spearman coefficient, which looks at rank order rather than linearity, also showed a strong correlation with a value of 0.87 (p < 0.0001) (Supplemental Table 1). Individual points are labeled with letters to indicate their amino acid replacements; wild-type is indicated as “WT”.

Based on the results presented in Figure 4, we performed a full kinetic analysis for select variants: wild-type, S267F (the NTCP*2 polymorphism), S267N, and S267W. Asparagine at position 267 was chosen because this substitution resulted in similar uptake as wild-type for estrone-3-sulfate and rosuvastatin but higher uptake for taurocholate. In contrast, the tryptophan substitution was chosen because it resulted in lower uptake for taurocholate but higher uptake for estrone-3-sulfate and rosuvastatin. For these four proteins, concentration dependent uptake was assessed under initial linear rate conditions using transiently transfected HEK293 cells. After normalizing for surface expression, we analyzed the results using the Michaelis-Menten equation and calculated K_m_ and V_max_ values (Figure 6, Table 1). Both the K_m_ and V_max_ were frequently altered, indicating altered substrate affinity and transporter turnover.

**Figure 6.**
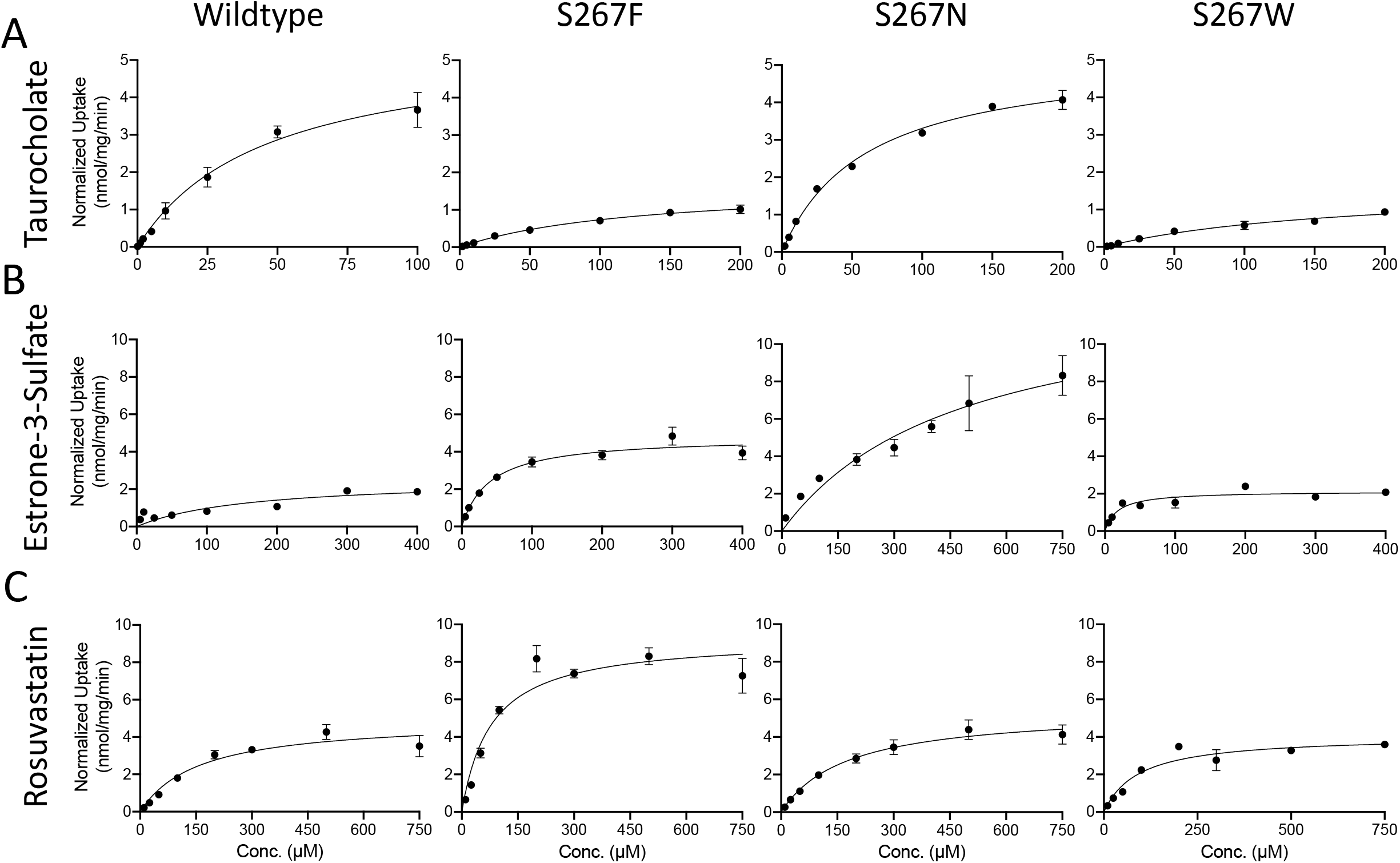
Kinetics of substrate transport mediated by wild-type NTCP and selected variants. Kinetics of taurocholate (A), estrone-3-sulfate (B), and rosuvastatin (C) uptake by wild-type NTCP (first column), S267F (second column), S267N (third column), and S267W (fourth column). Uptake of increasing concentrations of each substrate was measured under initial linear rate conditions in HEK293 cells 48 hours after transfection. Results shown are the mean ± SEM of a representative experiment completed with triplicate technical samples. The curves are best fits of the mean values using the Michaelis-Menten equation in GraphPad Prism 8. Results listed in Table report the average and SEM of at least 3 independent experiments, each comprising 3 technical replicates.

To summarize these data, we calculated the capacity of each variant to transport the various substrates (V_max_/K_m_) (Table 1). For the most part, the capacity of the variants chosen agreed with the rank order shown in Figure 4. Of the four variants, wild-type NTCP had the highest capacity (134 ± 14 μL/mg/min) for taurocholate (Figure 6A; Table 1) but the lowest capacity for the other two substrates: 15 ± 1.3 μL/mg/min for estrone-3-sulfate and 24 ± 5.1 μL/mg/min for rosuvastatin. The lowest capacity for taurocholate (11 ± 2.1 μL/mg/min) was determined to be for S267W (Figure 6A; Table 1) which is in agreement with the single time point single concentration results (Figure 4A). For estrone-3-sulfate and rosuvastatin (Figure 6B and C, Table 1), S267F showed the highest capacity with 99 ± 11 μL/mg/min and 112 ± 23 μL/mg/min, respectively, which confirms the results presented in Figure 4B and C. In summary, amino acid substitutions at rheostat position 267 differentially altered both kinetic parameters for substrate transport.

### Homology modeling of human NTCP structure

Both modeling and experiments suggest that human NTCP has 9 transmembrane (TM) helices (Doring, Lutteke, Geyer, & Petzinger, 2012; Hu et al., 2011) with an extracellular glycosylated amino terminus (Appelman, Chakraborty, Protzer, McKeating, & van de Graaf, 2017). These helices are arranged into “core” (TM 3, 4, 5, 8, 9 and 10) and “panel” domains (TM 2, 6 and 7) that flank the substrate-binding, intracellular crevice (*e.g*. Figure 1). Although no experimental structure is available for human NTCP, structural information could be derived from homology studies. Homologues in the SLC10A family exhibit 9.5 to 99.0% sequence identity and are present in most kingdoms of life (Supplemental Methods). Bacterial ASBTs are the best-structurally characterized; structures are available for *Yersinia frederiksenii* ASBT (ASBT_Yf_) (Zhou et al., 2014) and *Neisseria meningitidis* (ASBT_Nm_) (Hu et al., 2011).

In ASBT_Yf_, transport of bile acids and other substrates appears to be accomplished by a transition between inward-open and outward-open conformations. More specifically, this is accomplished *via* a rigid body motion of the core domain (Zhou et al., 2014) that allows alternating exposure of ligand-binding sites to the intracellular or the extracellular space. Both ASBT and NTCP co-transport sodium ions along with bile salts. In the crystal structure of ASBT_Nm_, two Na^+^ binding sites have been identified at the junction of core and panel domains (Zhou et al., 2014).

We used the inward-open and outward-open crystal structures of two bacterial ASBTs as templates for comparative modeling of human NTCP. We then applied all-atom structural refinement to the homology models to generate the lowest energy inward-open and outward-open homology models for wild-type human NTCP. In agreement with the bacterial structures, a well-defined pocket was present at the junction of the core and panel domains (Figures 1A and 1B), in which taurocholate could bind before being transported. Comparison of the inward- and outward-open models suggest that conformational changes in TM domains 3, 4a, 4b, 7, 9a, and 9b lead to closing of that pocket and an opening of a pocket where taurocholate binds before being released intracellularly (Figure 1C). Position S267 was located on TM9b, near the substrate binding cavity in both the inward- and outward-open conformations, and substitutions therefore may directly influence substrate transport.

### Evaluating stability changes arising from substitutions at position S267

Next, we modeled all 19 amino acid substitutions at position 267 of the wild-type homology models and assessed the predicted stability changes on the inward- and outward-open conformations. Starting with the inward-open conformation, we found that most substitutions were predicted to be stabilizing relative to the wild-type control (serine at position 267): 13 of the 19 potential substitutions yield energies more favorable (more negative) than serine (Figure 7A). This observation was striking because it is in stark contrast with typical results from such computation predictions, which have long found that the wild-type amino acid tends to score more favorably than any other substitution (Kuhlman & Baker, 2000).

**Figure 7:**
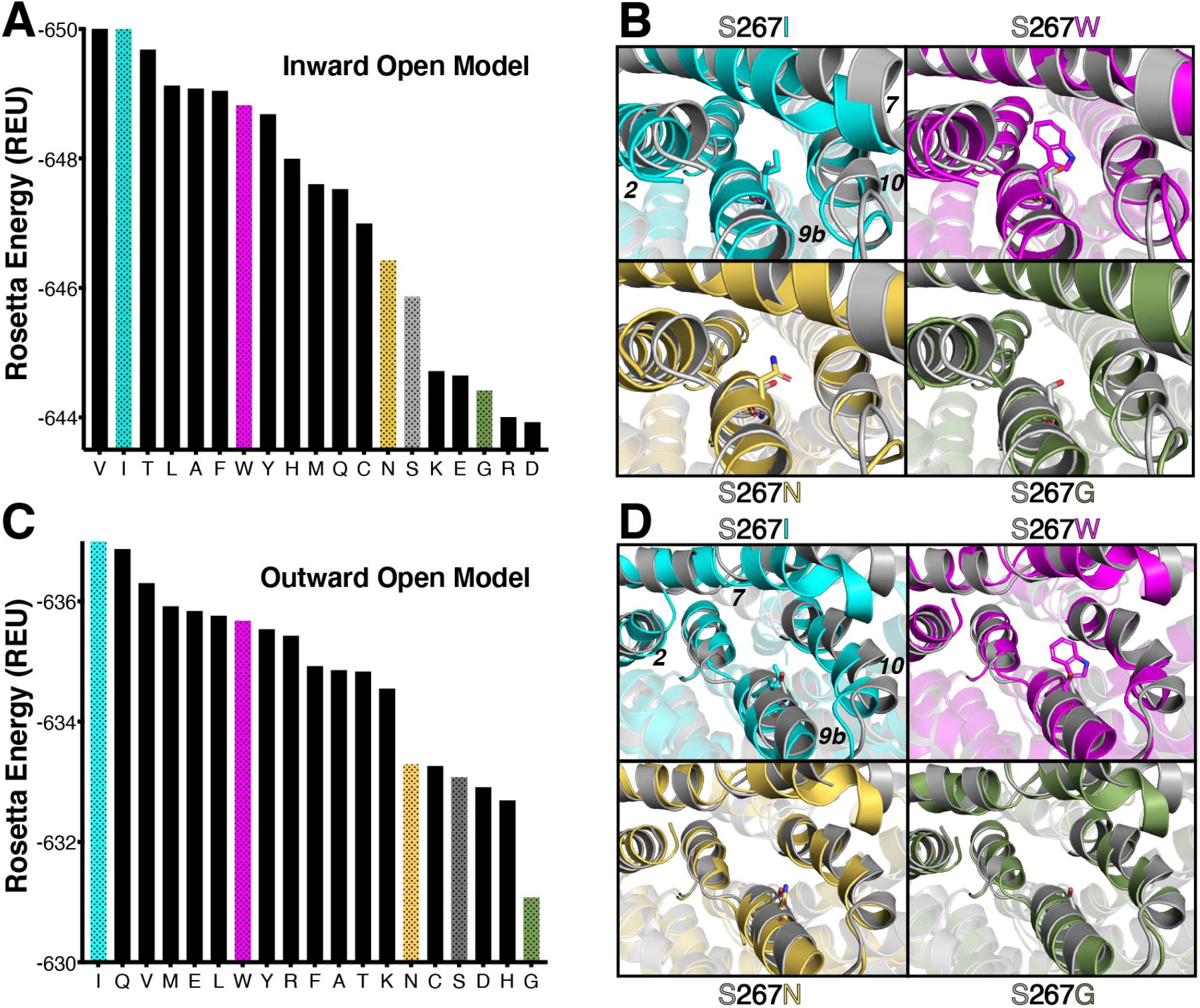
Predicted stability differences associated with sequence variation at the S267 position. (A) Rosetta energies for 19 sequence variants at the 267 position, using the inward-open model. Proline is not shown, because the energy associated with this residue is very unfavorable (off the scale); this is consistent with experimental results, in which proline showed the least amount of transport and somewhat diminished surface expression (Figures 1–3). (B) Structural details from the models underlying these energy differences. Four different sequence variants are compared with the wild-type S267; in each case, the conformation of TM helices 2 and 7 respond to changes in the amino acid at position 267, which is located on TM helix 9b. (C) Rosetta energies using the outward-open model. Proline is again not shown, because the energy associated with this residue is very unfavorable (off the scale). (D) Structural details from the outward-open models. In this conformation, the position of TM helix 10 responds to changes in the amino acid at position 267.

To explore the structural basis for the observed energetics in this modeling experiment, we selected two substitutions that were stabilizing (S267I and S267W), one with little effect on stability (S267N), and one predicted to be slightly destabilizing (S267G). Inspection of the representative conformations for each of these variants revealed slight changes in the local environment; in particular, TM helices 2 and 7 – which face the sidechain presented at position 267 – respond in a slightly different manner to each variant (Figure 7B).

In our model of the outward-open conformation, the wild-type S267 was more solvent exposed than in the inward-open conformation, with a solvent accessible surface area of 17.76 Å^2^ as compared to 4.26 Å^2^. Remarkably though, the same unusual behavior emerged when probing stability differences in the outward-open conformation. Many substitutions were predicted to be more stable than the wild-type serine (Figure 7C), and again the notable tolerance for alternate amino acids at position 267 could be rationalized by malleability of the local structure, this time the nearby TM helix 10 (Figure 7D). In both cases, the small rearrangement of these helices understates the dramatic differences in sidechain conformations needed to accommodate these alternatively packed arrangements (Supplemental Figure 1).

### Correlation of structure models and experimental data

We next compared the computational stabilities of each S267 variant to the experimental data. The most direct comparison should be to the NTCP surface expression, which would be decreased or increased by altered protein stability. To that end, we examined the effect between cellular expression levels and the calculated energies using (i) the inward-open NTCP model (Supplemental Figure 3A), (ii) the outward-open model (Supplemental Figure 3B), and (iii) the difference between the energies of the inward-open and the outward-open models (Figure 8). We anticipated that if the protein resides primarily in one conformation or the other, its surface expression may correlate with the calculated energies for that conformation; however, we find no statistically significant correlation between surface expression and the energies of the models in either conformation (Supplemental Table 1).

**Figure 8:**
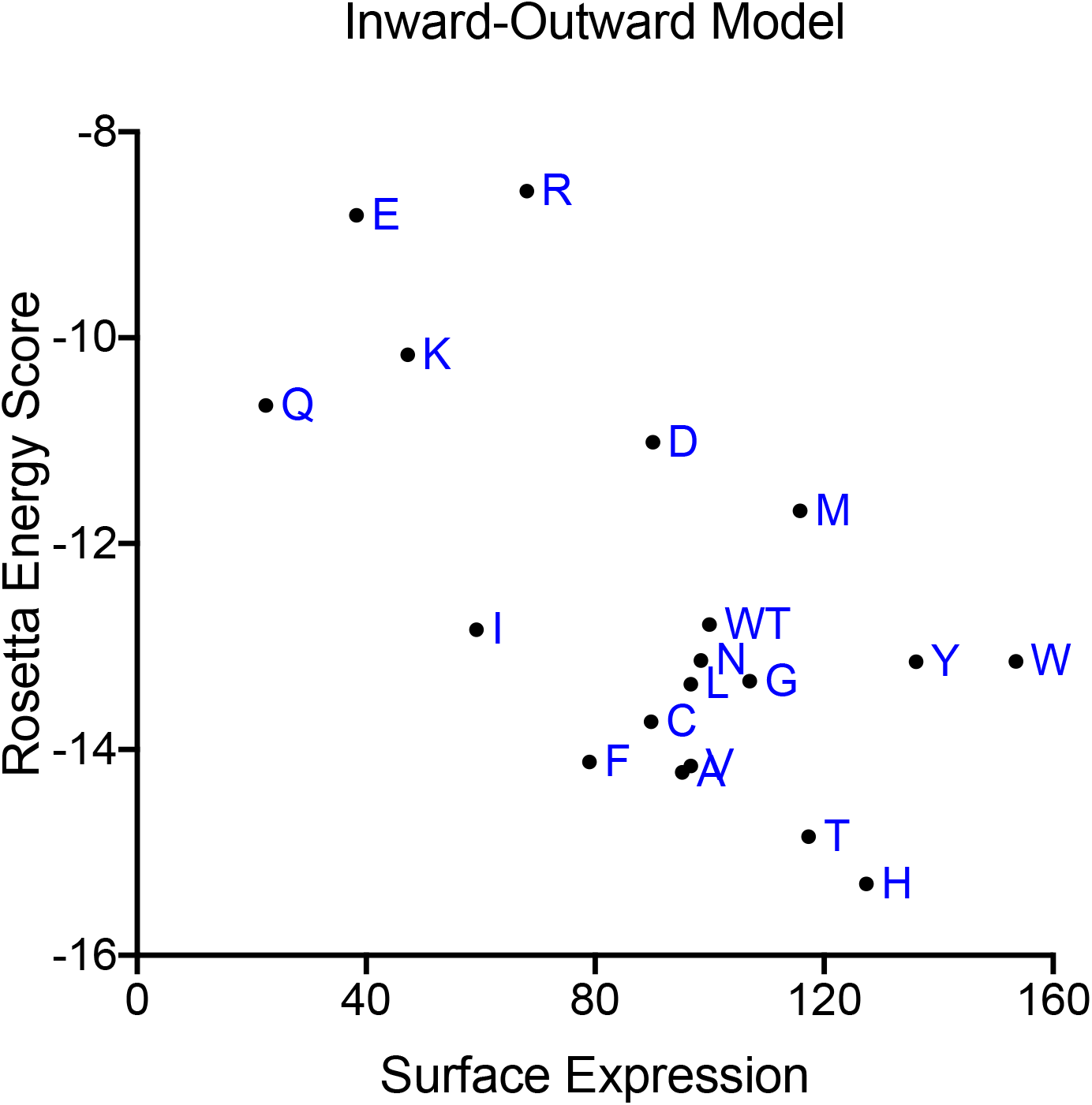
Correlation of Rosetta energy of the inward-open model minus energy of the outward-open model and quantification of surface expression levels. Energies calculated using Rosetta are plotted against the percent surface expression (Figure 2) of each variant. Individual points are labeled with letters to indicate their amino acid replacements; wild-type is indicated as “WT”. Proline is not shown because its predicted stability difference is off-scale.

Instead, we observed a correlation between surface expression and the difference in energy between the two conformations (Pearson and Spearman coefficients were −0.64 and −0.52, respectively; both correlation coefficients were non-zero with p < 0.05; Supplemental Table 1). There are several potential explanations for why sequence variants with more favorable energies in the inward-open model relative to the outward-open model (*i.e*., the difference between the states) yielded higher surface expression. A direct physical interpretation may be that the inward-open conformation promotes surface expression whereas a stable outward-open conformation serves as a “trap” that disfavors membrane insertion and/or correct folding required for presentation on the cell surface. In such a model, the outward-open conformation is critical for transport but over-stabilization of this conformation would prove deleterious. As an alternate explanation, however, we note that the Rosetta energy function is parameterized primarily for study of soluble proteins. As a result, energy differences between sequence variants are computed relative to a reference state that may not be appropriate for membrane proteins (the unfolded protein in a soluble context). Because of this, the outward-open conformation (in which position 267 is more solvent exposed) may simply be serving as an alternate (more appropriate) reference state, implying that calculated stability of the inward-open conformation is the primary determinant of surface expression, once energies have been more appropriately normalized.

For completeness, we additionally explored whether there was a relationship between the Rosetta-calculated energies and substrate uptake; we did not expect the Rosetta energies to be predictive, given that the balance of multiple conformational states is likely critical for effective transport. This expectation proved to be the case, as we did not observe a statistically significant correlation between any of the energies with update of any substrate (Supplemental Figure 2).

Overall, the strong correlation between surface expression and these computed energy differences supports the underlying structural rationale justifying how diverse amino acids can be accommodated at position 267 in a manner that altered – but did not disrupt – protein activity in a rheostatic manner. Since stability did not appear to be greatly altered, we hypothesize that the experimentally-observed changes in transport for each substitution arose from altered dynamics or from altered energies of other conformations that surely exist in the NTCP ensemble.

## Discussion

Rheostat positions were first described in soluble-globular proteins (Hodges et al., 2018; Meinhardt et al., 2013) and have been predicted in a wide variety of proteins (Miller, Vitale, Kahn, Rost, & Bromberg, 2019). However, to our knowledge, no such positions have been biochemically/experimentally identified in transmembrane proteins. In the present study, we confirmed that rheostat positions can occur in transmembrane transport proteins such as NTCP. Indeed, the natural polymorphic position, S267, behaved as a rheostat position for all three substrates tested: taurocholate, estrone-3-sulfate, and rosuvastatin (Figure 2).

The overall phenotype of each substitution at position 267 was primarily dominated by changes in altered transport kinetics. Full kinetic analyses with selected variants demonstrated that both K_m_ and V_max_ were differentially affected, along with substrate specificity. Nonetheless, both modeling results and experimental data indicated subtle changes in stability that were distinct from (not correlated with) changes in transport. Thus, changes at one rheostat position altered multiple functional and structural parameters, revealing a complex interplay that must be resolved to advance predictive pharmacogenomics. Since we also observed complex outcomes – affecting multiple functional parameters – arising from substitutions at rheostat positions in human liver pyruvate kinase, this complexity is likely common in a variety of proteins (Wu et al., 2019).

Of the affected functional parameters in NTCP, we were particularly intrigued by the altered substrate specificity that was demonstrated by the changed rank-orders (Figure 2, left side) and poor correlation (Figure 5) of amino acid substitutions for the three substrates. Historically, altered substrate specificity has been defined from the “perspective” of the proteins, as changes in either the rank-order of preferred substrate and/or changes in the fold-change of transport (Creighton, 1993; Tungtur et al., 2019). The results shown here – from the perspective of the substrate – provide an orthogonal view of specificity.

The substrate-dependent effects became even more apparent when we compared the kinetics for selected variants (Figure 6, Table 1). The substitution-dependent effects on substrate specificity could arise if the translocation pathway or binding pocket for taurocholate differed from that of the other two substrates. Distinct binding pockets within the translocation pathway were recently demonstrated for three substrates of the Organic Cation Transporter 1 (Boxberger, Hagenbuch, & Lampe, 2018). Different binding pockets or translocation pathways may be a common feature of multi-specific drug transporters. Alternatively, substrate-dependent substitution outcomes might arise if the different substrate/substitution combinations had different effects on the NTCP conformational and equilibrium dynamics, similar to the types of changes that arise from amino acid substitutions in beta-lactamase (Modi & Ozkan, 2018).

Indeed, substrate transport by Solute Carriers like NTCP is a dynamic process generally described by the ‘alternating-access’ model. As such, NTCP is an intrinsically flexible protein that undergoes complex and hierarchical conformational changes while carrying out its biological function of transporting substrates into hepatocytes. In addition to the outward- and inward-open conformations considered here, NTCP must have multiple intermediate states. One additional obligate conformation is an occluded state in which the binding site is not accessible from either side of the membrane. The stabilities and/or equilibrium dynamics for each of the conformations could be differentially affected by single amino acid substitutions, giving rise to the intermediate functional outcomes observed. Conformational changes are also modulated by interactions with substrates and sodium ions. Thus, an accurate evaluation of structural characteristics of intermediate conformations along the entire conformational transition pathway will be necessary to understand rheostatic substitution behavior. This remains a challenging task for both experimental and theoretical approaches.

When the rheostatic outcomes from the cellular uptake assays were further investigated, strong rheostatic effects were observed on transport, and the effects on stability of the inward-open conformation (in which position 267 was mostly buried) were slightly rheostatic. Furthermore, computational modeling and stability calculations were in agreement with the experimental measures of protein surface expression. This indicates that modeling will be a useful tool for identifying positions in functionally important regions that tolerate a wide range of substitutions, the hallmark characteristic of a rheostat position.

In conclusion, these combined results showed that NTCP position 267 is a rheostat position. Although individual substitutions had wide-ranging effects on the various aspects of function and stability that give rise to the overall phenotype of cellular transport, much of the change could be attributed to altered transport kinetics. Structurally, position 267 was strikingly tolerant to substitution, despite the fact that it was largely buried in the inward-open conformation. Based on these results, we propose that other polymorphic positions in NTCP and in other proteins might also be locations of rheostat positions (Fenton, Page, Spellman-Kruse, Hagenbuch, & Swint-Kruse, 2020). Given the complex interplay between substitutions and substrate specificity observed at this NTCP rheostat position, it is imperative to expand recognition and understanding of rheostat positions in order to advance predictive pharmacogenomics.

