## Supplemental for "A clinically-relevant polymorphism in the Na^+^/taurocholate cotransporting polypeptide (NTCP) occurs at a rheostat position"

**Supplemental Methods: Multiple Sequence Alignment**

A multiple sequence alignment of NTCP orthologs from a wide variety of species, including both prokaryotes and eukaryotes, was created using the PSI-BLAST (Basic Local Alignment Search Tool) algorithm (Altschul et al., 1997) from the National Center for Biotechnology Information. Eight separate searches were completed in July 2017. The first six searches were completed using the human SLC10A family members (Claro da Silva, Polli, & Swaan, 2013) as query sequences: NTCP (*SLC10A1*), ASBT (*SLC10A2*), P3 protein (*SLC10A3*), sodium/bile acid cotransporter 4 (*SLC10A4*), P5 (*SLC10A5*), and SOAT (*SLC10A6*). Several homologs were retrieved by two or more of the searches, which facilitated the later combination of these 6 alignments into one (below). The remaining BLAST searches were completed using two NTCP bacterial homologs, one from *Neisseria meningitidis* (pdb 3ZUY; (Hu, Iwata, Cameron, & Drew, 2011)) and one from *Yersinia frederiksenii* (pdb 4N7X; (Zhou et al., 2014)) as queries, each of which generated thousands of search hits with sequence identities spanning 39%-99%. Following each of the 8 BLAST searches, the resulting sequences were aligned using Clustal Omega (Madeira et al., 2019).

Next, to relate the eukaryotic and prokaryotic sequence alignments, the structural model built for human NTCP (see main text Methods) was super-imposed onto the 4N7X crystal structure of the bacterial homolog using UCSF Chimera (Pettersen et al., 2004). A structure-based sequence alignment was created from these reference proteins. This alignment was used in combination with PROMALS3D (Pei, Kim, & Grishin, 2008) to align the human SLC10A family members to the crystal structure. Finally, using sequences that were present in one or more alignments as guide sequences, the algorithm MARS (Parente, Ray, & Swint-Kruse, 2015) was used to combine the eukaryotic and prokaryotic sequences in one file without perturbing the structure-based alignments.

During these processes, sequences were curated by removing (i) duplicate sequences, (ii) particularly long or short sequences (as compared to human NTCP), (iii) sequences with large deletions, or (iv) those with ambivalent amino acid assignments. To prevent over-representation and facilitate proper sampling for subsequent calculations, bacterial sequences were clustering by their sequence identities; clear clusters were evident at the ~80% thresholds. Each of these clusters was randomly sampled to select no more than five sequences for inclusion in the final alignment of 1561 sequences (see sequence alignment file in the supplemental material) . Sequence entropy for the combined, sampled alignment was calculated using BioEdit v 7.0.5.3 software (Tippmann, 2004).

**Supplemental Table 1**: Pearson and Spearman coefficients calculated for various correlation studies.

| | Pearson | | Spearman | |
|---|---|---|---|---|
| | Value | p-value | Value | p-value |
| Taurocholate v Estrone-3-Sulfate | 0.38 | 0.1 | 0.39 | 0.09 |
| Taurocholate v Rosuvastatin | 0.33 | 0.16 | 0.23 | 0.33 |
| Estrone-3-Sulfate v Rosuvastatin | 0.82* | <0.0001 | 0.87* | <0.0001 |
| Taurocholate v Rosetta Inward | 0.054 | 0.83 | 0.16 | 0.52 |
| Estrone-3-Sulfate v Rosetta Inward | -0.14 | 0.58 | -0.12 | 0.63 |
| Rosuvastatin v Rosetta Inward | -0.15 | 0.54 | 0.075 | 0.76 |
| Taurocholate v Rosetta Outward | -0.11 | 0.66 | 0.28 | 0.24 |
| Estrone-3-Sulfate v Rosetta Outward | 0.065 | 0.79 | 0.18 | 0.46 |
| Rosuvastatin v Rosetta Outward | 0.066 | 0.79 | 0.25 | 0.3 |
| Taurocholate v Rosetta Inward Minus Outward | 0.15 | 0.53 | -0.074 | 0.76 |
| Estrone-3-Sulfate v Rosetta Inward Minus Outward | -0.21 | 0.39 | -0.3 | 0.21 |
| Rosuvastatin v Rosetta inward Minus Outward | -0.23 | 0.35 | -0.17 | 0.48 |
| Surface Expression v Rosetta Inward | -0.32 | 0.18 | -0.25 | 0.31 |
| Surface Expression v Rosetta Outward | 0.32 | 0.18 | 0.3 | 0.21 |
| Surface Expression v Inward Minus Outward | -0.64* | 0.0034 | -0.52* | 0.023 |

Correlation scores from comparison figures (Figures 5, 8, and Supplemental Figure 3) were calculated in GraphPad Prism 8 using either the Pearson correlation or the nonparametric Spearman correlation.

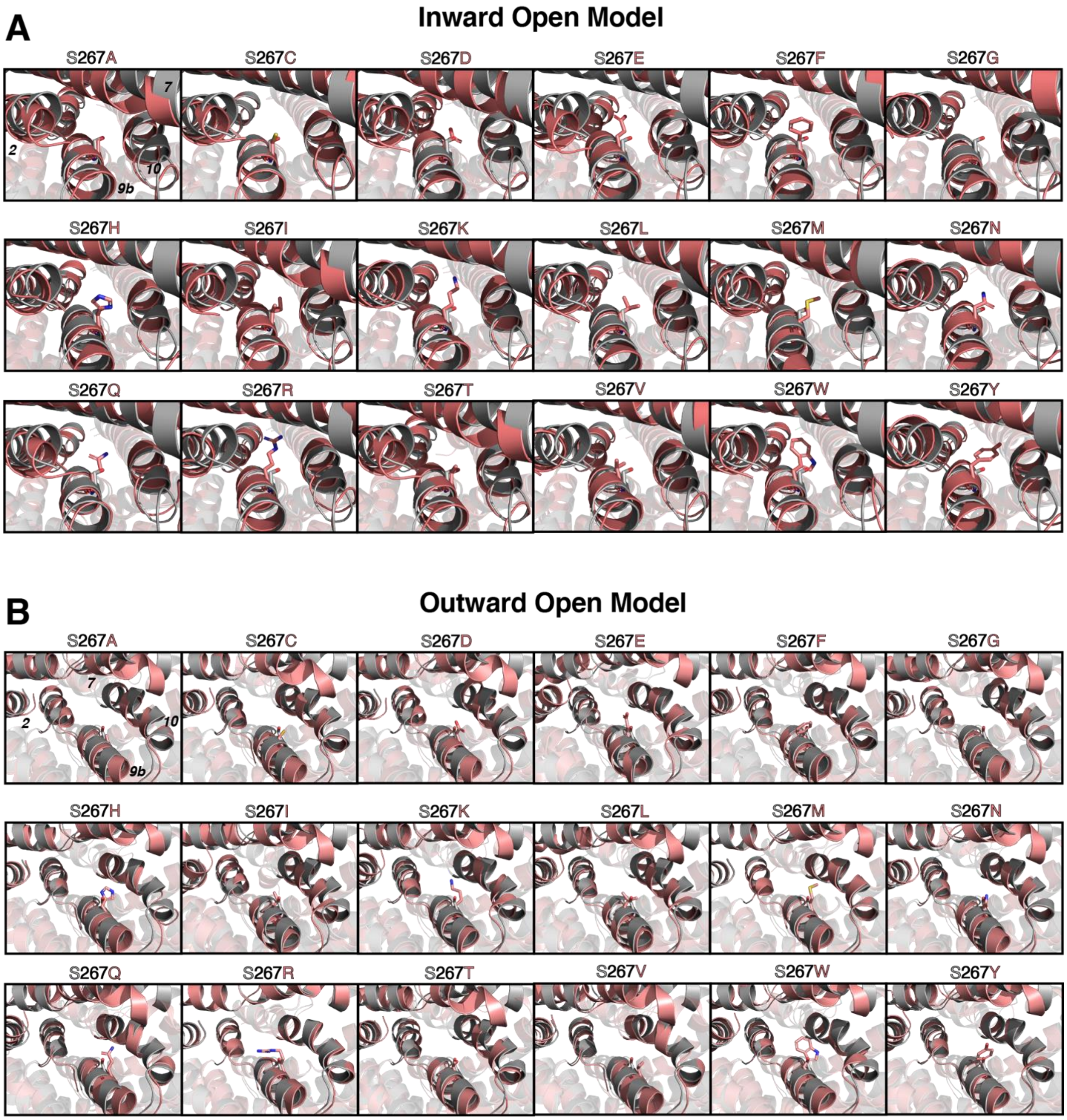


**Supplemental Figure 1: Sequence variation at S267 position leads to alternative local packing.** Structural details are shown from WT in gray and 18 mutant structures in salmon color (proline was excluded) at the 267 position. (A) Models in the inward-open conformation. (B) Models in the outward-open conformation.

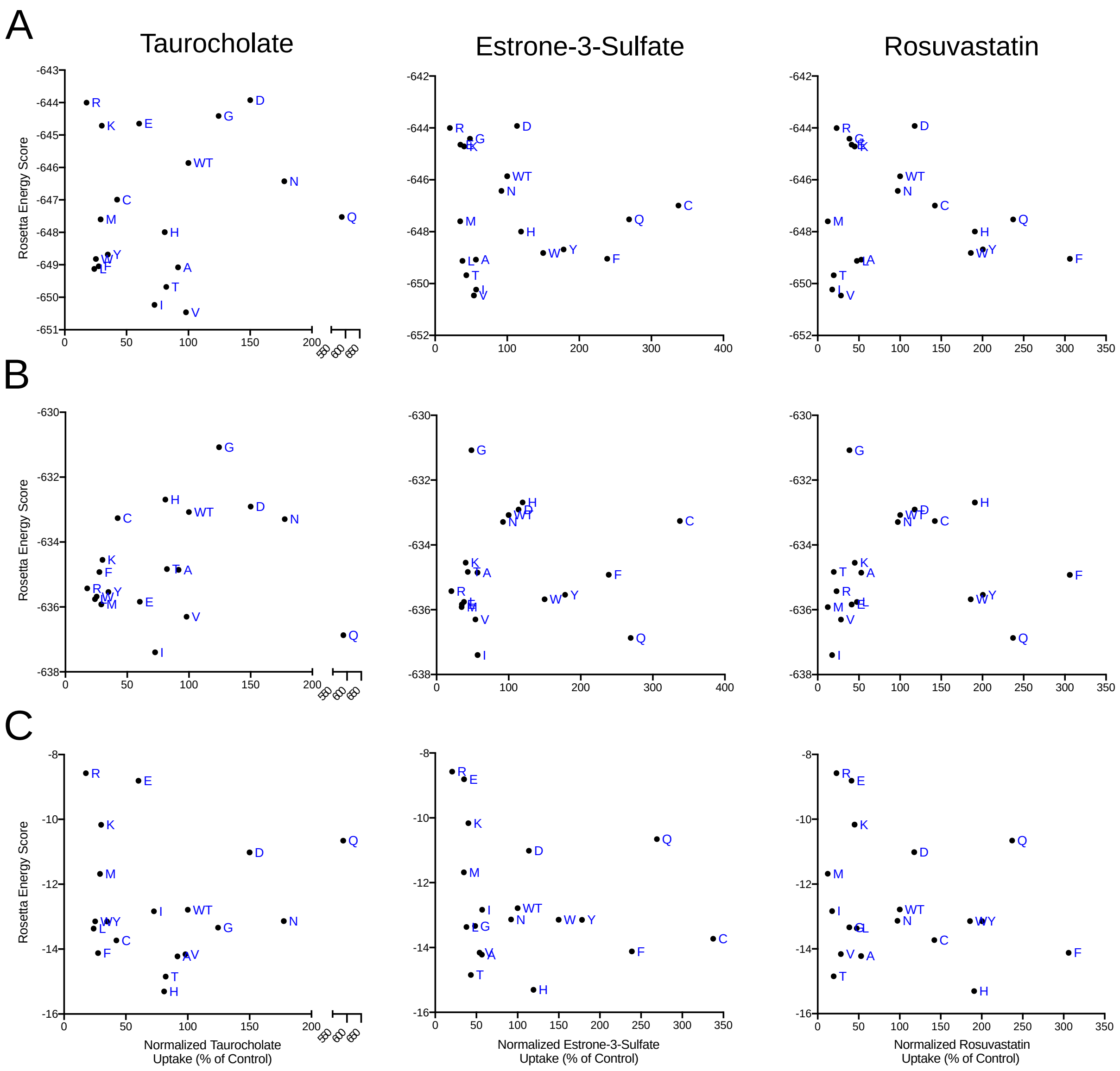


**Supplemental Figure 2: Correlation of Rosetta energy scores and normalized initial substrate uptake.** Inward-open model (A), outward-open model (B) and inward-open minus outward-open model (C) energy scores calculated using the Rosetta software suite are plotted against normalized substrate uptake values from Figure 3. No correlations were observed.

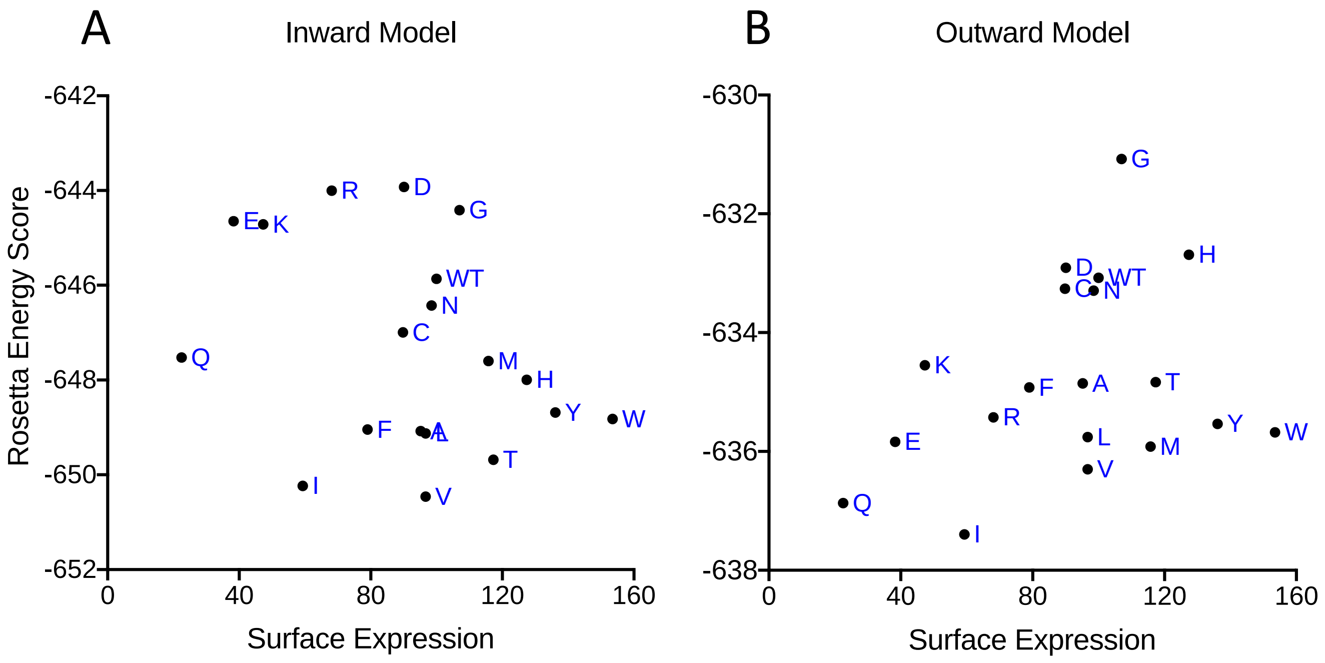


**Supplemental Figure 3: Correlation of Rosetta energies and quantification of surface expression levels**. Energies calculated using Rosetta are plotted against the percent surface expression (Figure 2) of each variant. Individual points are labeled with letters to indicate their amino acid replacements; wild-type is indicated as “WT”. (A) Energy for the inward-open model, (B) energy for the outward-open model
